# Rapid and objective assessment of auditory temporal processing using dynamic amplitude-modulated stimuli

**DOI:** 10.1101/2024.01.28.577641

**Authors:** Satyabrata Parida, Kimberly Yurasits, Victoria E. Cancel, Maggie E. Zink, Claire Mitchell, Meredith C. Ziliak, Audrey V. Harrison, Edward L. Bartlett, Aravindakshan Parthasarathy

## Abstract

Auditory neural coding of speech-relevant temporal cues can be noninvasively probed using envelope following responses (EFRs), neural ensemble responses phase-locked to the stimulus amplitude envelope. EFRs emphasize different neural generators, such as the auditory brainstem or auditory cortex, by altering the temporal modulation rate of the stimulus. EFRs can be an important diagnostic tool to assess auditory neural coding deficits that go beyond traditional audiometric estimations. Existing approaches to measure EFRs use discrete amplitude modulated (AM) tones of varying modulation frequencies, which is time consuming and inefficient, impeding clinical translation. Here we present a faster and more efficient framework to measure EFRs across a range of AM frequencies using stimuli that dynamically vary in modulation rates, combined with spectrally specific analyses that offer optimal spectrotemporal resolution. EFRs obtained from several species (humans, Mongolian gerbils, Fischer-344 rats, and Cba/CaJ mice) showed robust, high-SNR tracking of dynamic AM trajectories (up to 800Hz in humans, and 1.4 kHz in rodents), with a fivefold decrease in recording time and thirtyfold increase in spectrotemporal resolution. EFR amplitudes between dynamic AM stimuli and traditional discrete AM tokens within the same subjects were highly correlated (94% variance explained) across species. Hence, we establish a time-efficient and spectrally specific approach to measure EFRs. These results could yield novel clinical diagnostics for precision audiology approaches by enabling rapid, objective assessment of temporal processing along the entire auditory neuraxis.

## Introduction

Perception of speech and other natural sounds critically depends on the auditory system’s ability to encode temporal cues, such as the envelope and fine structure, which convey complementary information present in sounds (Rosen, 1992; Smith et al., 2002). Neural coding deficits of either cue can result in sustained auditory perceptual deficits (Moore, 2019). Current clinical hearing tests, e.g., pure-tone audiograms, are not designed to capture these suprathreshold neural coding deficits, rather focusing primarily on near-threshold cochlear function and hearing sensitivity (Moore, 2012; Ruggles et al., 2011; Strelcyk & Dau, 2009). Approximately one in ten patients seeking hearing health care have complaints of hearing in noise, despite normal audiometric thresholds (Cancel et al., 2023; Parthasarathy et al., 2020; Spehar & Lichtenhan, 2018; Tremblay et al., 2015). These hearing deficits, which remain unaddressed by current clinical measures, represent a large unmet medical need, and necessitate the development of sensitive and objective clinical tests of hearing function.

Auditory evoked potentials may facilitate objective neural assessment of auditory processing. The auditory brainstem response (ABR) driven by brief, discrete sounds, are primarily used to assess hearing thresholds when the patient cannot behaviorally respond, such as in the newborn hearing screening or during skull-base surgery (Eggermont, 2019; Parkkonen et al., 2009). FFRs elicited by short speech tokens show more promise in assessing suprathreshold coding of envelope and fine structure cues, but these complex-FFRs superpose neural generators due to smear from temporally overlapping responses (Coffey et al., 2019). An ideal middle-ground may be provided by the envelope following response (EFR) which represents the polarity-tolerant (carrier-insensitive) component of FFRs and is elicited by the envelope of simpler amplitude-modulated (AM) carriers (Kuwada et al., 2002; Parthasarathy & Bartlett, 2011). The ability of neurons to phase-lock to the temporal envelope is biophysically constrained, such that while the auditory nerve can phase-lock to rates > 1500 Hz, neurons in the auditory cortex can phase-lock up to ∼80 Hz (Joris et al., 2004). Thus, by changing the characteristics of the amplitude envelope, we can emphasize different generators (Herdman et al., 2002; Kuwada et al., 2002; Parthasarathy & Bartlett, 2012; Parthasarathy & Kujawa, 2018). The EFR has been used as an objective neural correlate of perceptual performances in normal-hearing populations, for example, to explain individual variability in AM perception or the effect of language or musical training on temporal coding and perception (Krishnan et al., 2005; Lee et al., 2009; Purcell et al., 2004). In addition, the subcortical versus cortical signatures of AM vary across hearing loss etiologies - the EFR has been used to understand AM coding changes due to age-related (Dimitrijevic et al., 2016; Parthasarathy et al., 2016a, 2019; Parthasarathy & Bartlett, 2011), permanent (Ananthakrishnan et al., 2016; Han et al., 2021; Parida & Heinz, 2021; Race et al., 2017), and noise-induced ‘hidden’ (Shaheen et al., 2015; Wilson et al., 2021) hearing loss etiologies.

Unfortunately, the utility of EFRs as a clinical tool is limited because current practices typically involve collecting EFRs serially for each AM frequency, which is time consuming. Previous attempts to speed up data collection involved multiplexing multiple carrier- and AM-frequency combinations (Dolphin, 1996; Holmes et al., 2018; John et al., 1998; Varghese et al., 2015) or multiband harmonic complexes (with different AM frequency in different carrier bands) (Dolphin, 1996; Wang et al., 2019). However, EFRs responses to such simultaneous stimulus presentations can be affected by across-frequency auditory processing and other nonlinearities, which are irrelevant to AM processing (John et al., 1998; Parthasarathy et al., 2016b; Wang et al., 2019). Discrete EFR tokens may also be confounded by heterogeneity in head size and neural generator locations that may result in subject-specific effects on particular AM frequencies (Shaheen et al., 2015).

Here we develop a new approach to efficiently assess neural AM coding using EFRs and validate it in four different species (humans, gerbils, rats, and mice). Our approach uses a dynamically varying AM (dAM) stimulus in combination with spectrally specific analyses, which yields robust high-SNR EFRs. The dAM stimulus varies continuously across AM frequencies (e.g., 16-1500 Hz over 1 s) and therefore allows for a richer estimation of EFRs with continuously sampled AM frequencies within a specified range. Estimating EFRs using the dAM stimulus is time efficient (approximately 5× faster) compared to sequentially sampling AM frequencies even when using half-octave wide steps. These dAM-stimuli-elicited EFRs are analyzed using spectrally specific tools (Parida et al., 2021), which provide optimal spectro-temporal resolution compared to traditional signal-processing metrics such as windowed fast Fourier transforms. The resultant EFR-metrics are highly similar to traditional EFR-metrics derived from the same subjects using discrete (stationary) SAM stimuli. These results highlight the efficacy of dAM-stimuli-derived EFRs to assess temporal coding by the auditory system and open up the possibility of utilizing EFRs in the clinic as a rapid assessment of supra-threshold auditory temporal processing.

## Results

### Dynamically amplitude modulated stimuli are time efficient to probe EFRs to multiple modulation rates

Neural phase-locking to the stimulus amplitude envelope has been typically probed using discrete sinusoidally amplitude modulated (SAM) tones, where both the modulator (*f*_*m*_) and the carrier (*f*_*c*_) are sinusoids (i.e., frequencies are stationary). In contrast, we introduce a dynamically amplitude modulated stimulus, where the carrier is constant, but the modulation frequency varies exponentially, e.g., from 16 Hz to 1500 Hz (Fig 1). We have adopted these values based on subcortical and cortical tuning to amplitude modulation (see *Materials and Methods*). Using such dynamic AM trajectories, the AM coding by the auditory system can be studied in a time-efficient manner. For example, using a 3.33 Hz presentation rate, 200 trials per AM frequency, and 1/2 octave steps, it takes approximately 15 minutes to complete testing from 16-2048 Hz, similar to previous protocols (e.g. Parthasarathy & Bartlett, 2012). In order to cover a comparable range from 16 to 1500 Hz used in the current study, it takes approximately 3 min. Thus, the dAM stimulus offers 5x improvement in time efficiency over conventional SAM-tone-based approaches.

**Fig 1.**
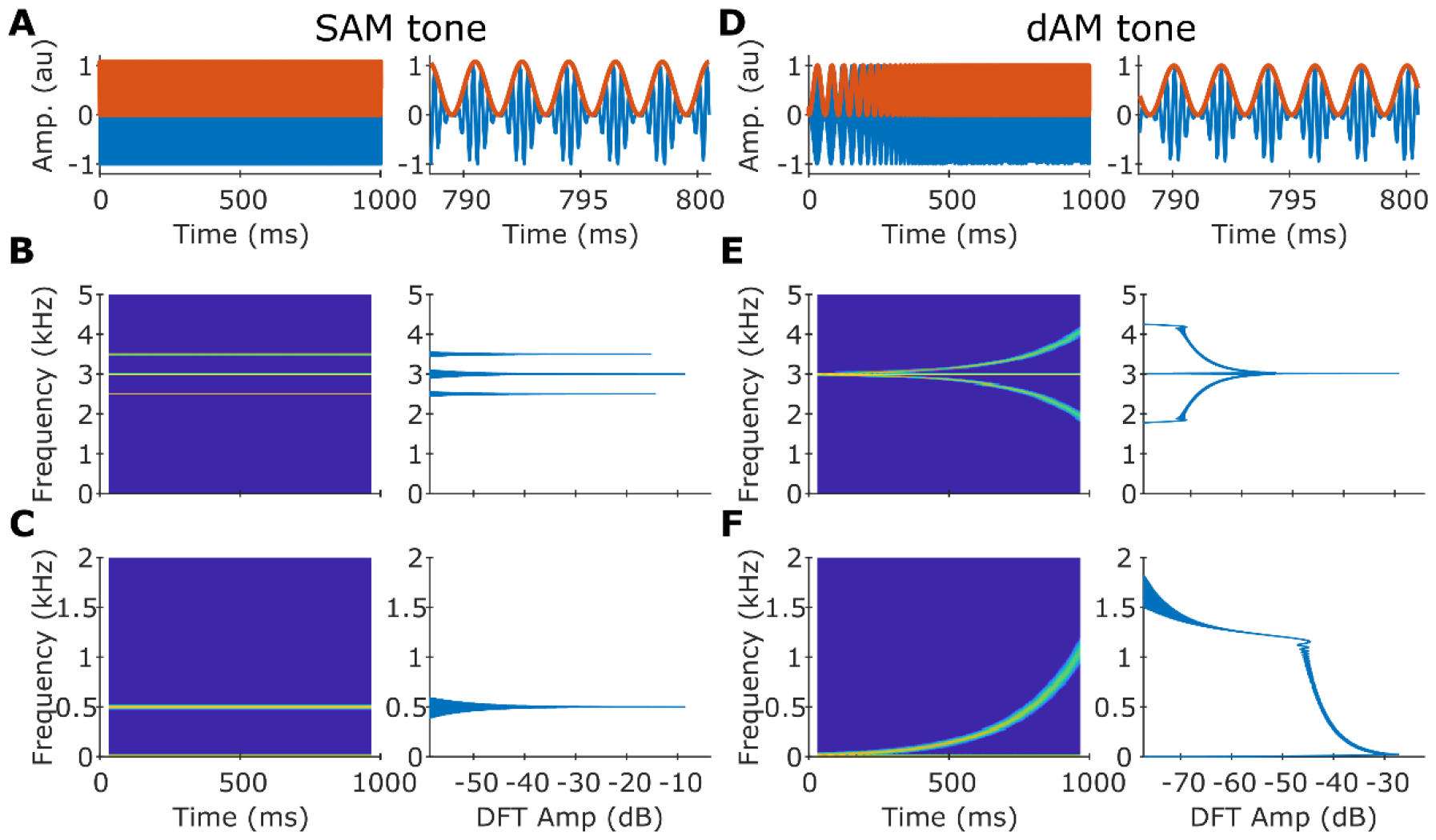
Spectrotemporal representations of a sinusoidally amplitude modulated (SAM) tone and a dynamically amplitude modulated (dAM) tone. (A) Time-domain waveforms of the SAM tone (blue) and its Hilbert envelope (red) over 1 s and when zoomed in over 15 ms. The SAM tone had a carrier of 3 kHz, and it was modulated with a 500 Hz AM frequency at 100% depth. (B) The spectrogram (left) and the discrete Fourier transform (DFT; right) magnitude of the SAM tone. (C) Same format as B but for the demeaned Hilbert envelope of the SAM tone. (D-F) Same format as A-C but for the dAM tone (carrier = 3 kHz, exponentially increasing AM from 16 Hz to ∼1200 Hz over 1 s). The dAM tone had an AM frequency of 500 Hz around 795 ms. Spectrograms were constructed using a 64-ms Hamming window with 95% overlap.

### Spectrally specific analyses offer optimal spectrotemporal resolution compared to traditional signal processing methods

Traditionally, time-varying signals are analyzed using a windowing method, such as the spectrogram. In each window, the signal is windowed using a taper (e.g., Hamming) and a power spectral density is estimated. The duration of the window determines the spectral resolution of the power spectral density. For example, for a 25-ms window, the spectral resolution is 40 Hz (inverse of the window duration). However, a longer window does not always offer better spectral resolution for dynamic signals because spectral trajectory can dramatically vary for longer windows. We quantified these time-frequency uncertainty effects as the time-bandwidth product (which denotes the overall spectrotemporal resolution) required to capture 95% of the dAM stimulus for various window durations. Specifically, for a given window duration, we divided the stimulus into nonoverlapping segments (two examples in Fig. 2A-B). For each segment, we estimated the signal’s power spectral density (after multiplying with a hamming window) and estimated the frequency bandwidth that captured 95% of the windowed-signal power. We define the product of the window duration and the frequency bandwidth as the spectrotemporal resolution (unitless) for that segment. The sum of this time-bandwidth product over all segments is the spectrotemporal resolution of the signal for the given window duration. For the dAM stimulus employed in this study and for windows ranging from 10 ms and 200 ms, the spectrotemporal resolution showed a U shape with values ranging between 86.4 and 255, with the best (minimum) spectrotemporal resolution occurring at 50 ms (Fig. 2C).

**Fig 2.**
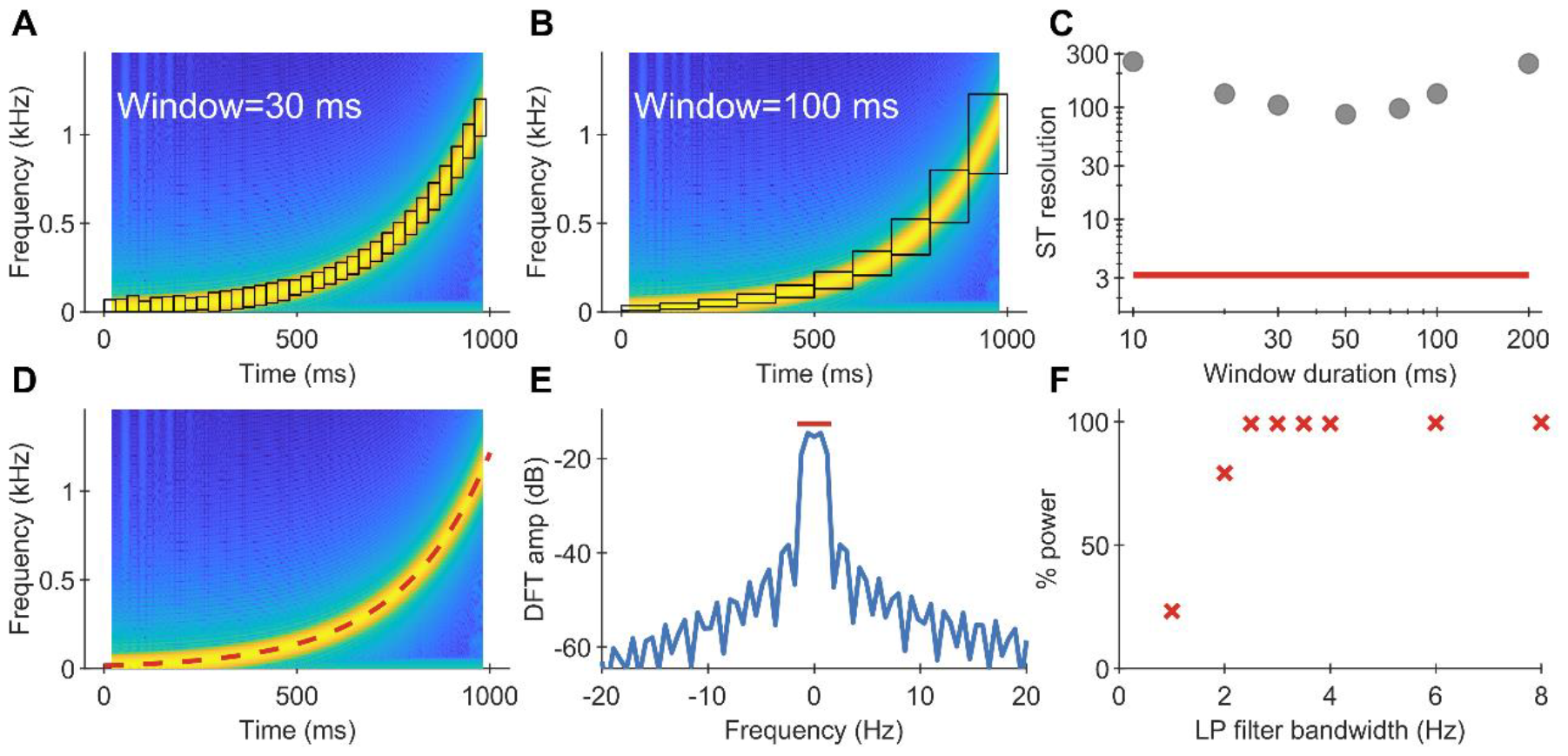
Spectrally specific analyses offer optimal spectrotemporal resolution. (A-B) The spectrogram of the demeaned Hilbert envelope of the dAM tone. Black boxes indicate the spectrotemporal tiles required to capture at least 95% of the envelope power of this dynamic stimulus for 30 ms (A) and 100 ms (B) analysis-window durations. The cumulative area of these red boxes is defined as the spectrotemporal resolution for a given analysis-window duration. (C) Spectrotemporal resolution for different analysis-window durations (gray symbols) and for spectrally specific analysis (red line), as described next. (D) The dAM trajectory (red dashed line) overlayed on the envelope spectrogram. (E) The DFT magnitude of the frequency-demodulated dAM envelope. The red line indicates the spectral window (∼3 Hz) required to capture ∼95% of the envelope power. (F) Percentage power captured of the dAM-tone envelope as a function of spectral bandwidth around 0 Hz. Spectrogram dynamic range = 60 dB in all panels.

However, spectrotemporal resolution can be greatly improved by using spectrally specific analyses compared to these windowing methods. Specifically, spectrally specific analyses utilize the knowledge of the spectral trajectory using other signal processing techniques (such as the Hilbert transform, frequency demodulation, and low-pass filtering) to track the power in a mathematically optimal window along the trajectory (Fig 2D). To highlight the improvement in spectrotemporal resolution, we estimated the discrete Fourier transform of the frequency demodulated dAM stimulus (Fig 2E) and estimated the total signal power captured as a function of low-pass filter bandwidth. This function saturated quickly around 3 Hz for the 1-s long signal, which was sufficient to capture >95% of the total stimulus power (Fig 2F). Thus, for the dAM stimulus used in our study, spectrally specific analyses offer ∼30x improvement in spectrotemporal resolution over windowing-based approaches. This superior resolution will improve the SNR of the power metric as power is integrated over a narrow spectrotemporal region that corresponds to the desired dAM trajectory, thus reducing the effects of unwanted noise.

### The temporal modulation transfer function (tMTF) can be efficiently estimated by combining the dAM stimuli and spectrally specific analyses

A key focus of the study was to (time) efficiently derive EFR amplitudes as a function of varying amplitude modulation rates, also referred to as the temporal modulation transfer function (tMTF). Traditionally, tMTFs for EFRs using discrete AM tokens are estimated by performing spectral FFT analysis on individual responses (Fig 3A). For example, each dashed box corresponds to a different AM frequency, with five predefined AM frequencies in total. In contrast, as we describe next, the tMTF can be efficiently estimated by combining the dAM stimulus and spectrally specific analyses to obtain a denser frequency sampling (Fig 3B). The EFR in response to the dAM stimulus tracks the dAM trajectory, the power along which can be extracted using spectrally specific analysis (i.e., Hilbert, frequency demodulation, and low-pass filtering). The low-pass filter can be implemented in the time domain as well as the spectral domain depending on what it desired. Time-domain filtering (e.g., using a Butterworth filter) enables one to track trajectory power on a sample-by-sample basis (i.e., at the sampling frequency of the signal without the need for binning (Fig 3B). However, time-domain filters use a single impulse response, estimating power along the trajectory using the variance of the signal and can be prone to bias and variance (Babadi & Brown, 2014; Parida et al., 2021). Since the relationship between dAM trajectory and time is known (which, in our case, is one-to-one), the output can be represented as dAM-trajectory power as a function of dAM-trajectory frequency. This function is a continuously sampled tMTF. In contrast, low-pass filtering can be implemented in the frequency domain on the multi-taper spectral estimates that help reduce bias and variance and are preferred if overall power is desired in longer windows. To estimate a tMTF that is analogous to the traditional discretely sampled tMTF (say at f1 Hz AM frequency), one could simply window this continuously sampled tMTF, where the (time) window is restricted to the vicinity of the f1 AM frequency (e.g., a 50-ms window).

**Fig 3.**
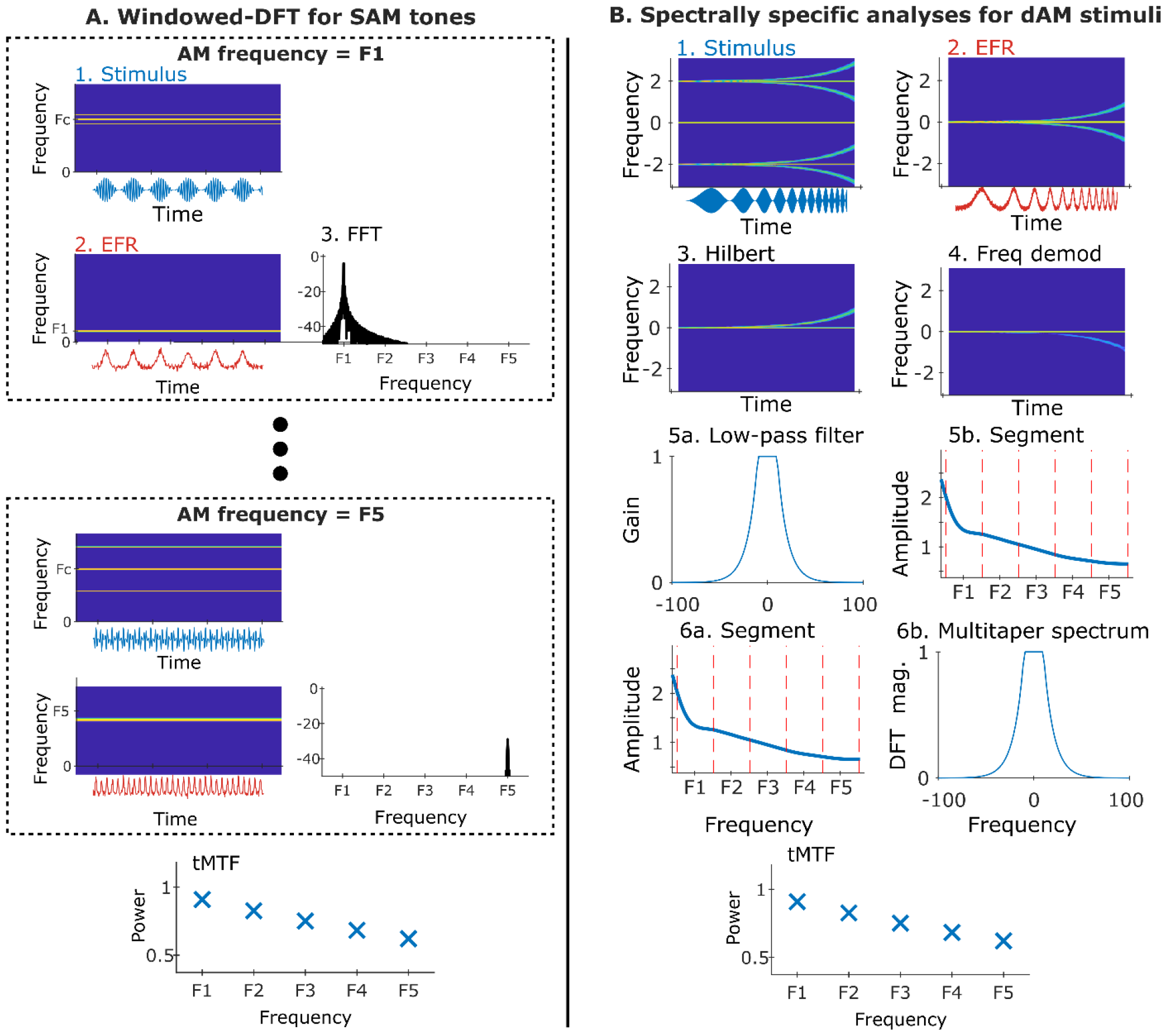
Schematic illustrating traditional and proposed methods to estimate the tMTF. (A) Traditionally, for a target AM frequency (e.g., F1), EFRs (several trials of opposite polarities) are collected in response to the corresponding SAM tone. The modulation coding strength at F1 (i.e., the tMTF value at F1) is defined as the DFT magnitude of the EFR at F1. This procedure is repeated for all target AM frequencies (F1-F5). (B) Our proposed method involves a single dAM stimulus, in response to which EFRs of opposite polarities are collected. Next, we compute the demeaned Hilbert envelope of the EFR, and frequency demodulate it using the known dAM trajectory. To estimate the tMTF value at an AM frequency (e.g., F1), first we choose a time window where the dAM trajectory is within a certain bandwidth (e.g., 0.2 octave). Note that since the relationship between AM frequency and time is one-to-one, the x-axis can have time or AM frequency as the unit. The tMTF value in this window can be estimated two ways: i) by low-pass filtering the frequency-demodulated signal in the time domain and taking the variance (=power) of the signal within the appropriate window, or ii) windowing the frequency-demodulated signal and estimating the power in low-pass window from its multi-taper spectrum.

### Robust EFR to dAM stimuli can be obtained for various species/carrier types

We recorded EFRs in response to the dAM stimuli in humans, gerbils, rats, and mice (Fig 4). Stimulus envelope features were noticeably captured in the dAM EFR grand averages (i.e., average EFR across all subjects in individual species) both in the time-domain waveform (e.g., low-frequency oscillations before 500 ms in mice and rats) as well as in the spectrograms (multiple harmonics of the dAM frequency trajectory). Note that EFRs in rats were recorded in a two-channel configuration similar to prior studies (Parthasarathy & Bartlett, 2012; Race et al., 2017), with channel 1 emphasizing peripheral generators, and channel 2 emphasizing central auditory generators, in response to both tone (not shown) and noise carriers.

**Fig 4:**
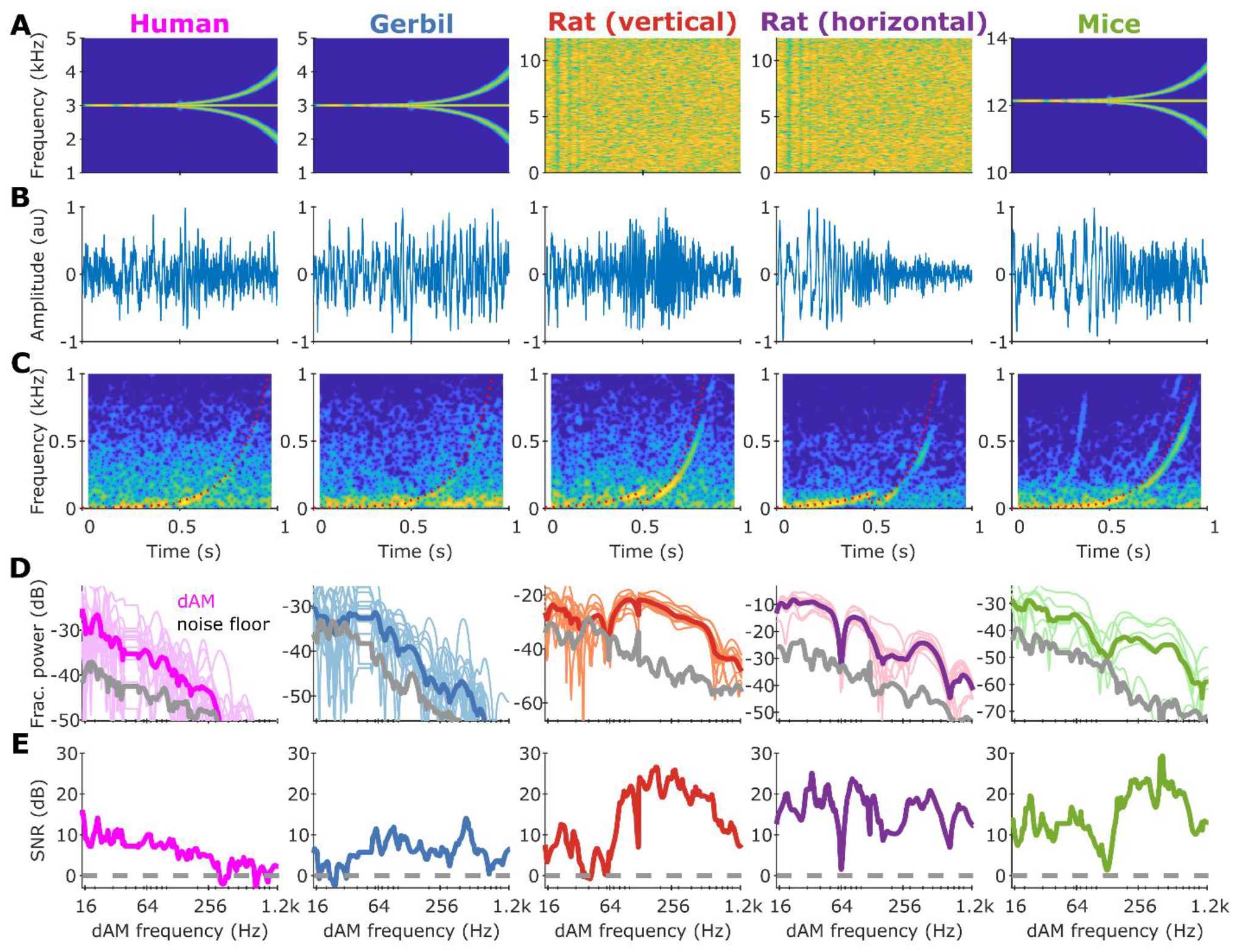
Robust EFR to dAM stimuli can be obtained for various species/carrier types. (A) The stimuli used to obtain the EFR response in several species (columns). The carrier frequency was species specific. For rats, both tone (not shown) and noise carriers were used in two different recording configurations that emphasize more subcortical (vertical) or cortical (horizontal) contributions. (B) Time-domain grand average EFR waveforms (i.e., averaged across all subjects within a species). (C) Spectrograms of the grand-average EFR responses with the dAM trajectory overlaid (red dotted lines). For all species, the second harmonic of the dAM trajectory is also evident. (D) Tracked fractional power (see Methods) along the dAM trajectory (colored) and along the noise floor trajectory (gray, parallel to the dAM trajectory but shifted up by 36 Hz) within a 24 Hz bandwidth. (E) Signal-to-noise ratio estimated using the average traces in D. EFR responses for all species, especially rats and mice, generally showed robust high-SNR responses for AM frequencies between 16 Hz and 1200 Hz.

We quantified the power along the dAM trajectory using spectrally specific analyses, as described before. We also estimated a noise floor for each species using an AM trajectory that was parallel shifted up by 36 Hz to the dAM trajectory (see *Materials and Methods*). Tracked power along the dAM trajectory was substantially above the noise floor across all species, although the exact values varied across frequency and species. dAM power in general showed a low-pass shape, which is typical of EFR responses (Lai et al., 2017; Parthasarathy & Bartlett, 2011). Next, we quantified the signal-to-noise ratio of the tracked dAM power (i.e., tracked dAM power minus the time-varying noise floor). Human-scalp recorded EFRs showed ∼10 dB SNR below 100 Hz and fell consistently below the noise floor above 800 Hz. These values are consistent with previous EFR studies in humans (He et al., 2008; Wiinberg et al., 2019).

All rodent species showed much higher SNRs, especially at higher AM frequencies. EFRs for all rodent species were well above noise floor even at 1400 Hz AM frequency. Interestingly, compared to rat EFRS recorded in the vertical electrode configuration (i.e., forehead to ipsilateral mastoid), rat EFRs recorded in the horizontal configuration (i.e., interaural line to ipsilateral mastoid) had better SNR at low AM frequencies and poorer SNR at high AM frequencies, consistent with previously published studies (Lai et al., 2017; Parthasarathy & Bartlett, 2012; Race et al., 2017). The horizontal configuration is thought to receive substantial contributions from midbrain and forebrain sources, which are sensitive to low-AM frequencies, while the vertical configuration is thought to emphasize neural generators from the peripheral auditory pathway. Hence, trends previously demonstrated using EFRs to discrete AM tokens are evident in the dAM EFRs as well. In summary, the dAM stimulus elicits robust, high-SNR EFRs across a wide-range of AM frequencies, and these EFRs can distinguish complementary neural generators (e.g., subcortical vs cortical) along the auditory neuraxis.

### dAM-stimuli-derived tMTFs are highly correlated with SAM-stimuli-derived tMTFs

Next, to validate our method to estimate the tMTF, we compared dAM-derived tMTFs to traditional tMTFs (i.e., tMTFs derived using discrete SAM tones) in the same group of subjects across all species. The number of AM frequencies in the traditional tMTF were different across species due to experimental time constraints (e.g., four AM frequencies for humans as data collection takes longer per frequency). The tMTF value at each AM frequency was the DFT magnitude of the EFR response for the corresponding condition. These tMTFs showed the expected low-pass shape. Additionally, similar to the continuously sampled dAM tMTFs for rats, the SAM tMTFs for rats showed higher magnitude at low (<100 Hz) frequencies for the horizontal configuration (compared to the vertical configuration) and lower magnitude at higher (>100 Hz) frequencies.

To facilitate direct comparison between traditional tMTFs and dAM-derived tMTFs, we constructed a discretely sampled tMTF at the same AM frequencies as in traditional TMTFs by windowing the frequency-demodulated dAM EFR (see Fig. 3, *Material and Methods* for details) centered at when the dAM trajectory crossed the AM frequency, estimated its multi-taper spectrum, and computed power in a low-frequency band. These tMTFs were similar to the traditional tMTFs for all species and electrode configurations tested (Fig. 5A-E) and were significantly correlated individually for each species as well as when pooled together (Fig. 5F). In summary, these results validate our approach of efficiently estimating the tMTF.

**Fig 5:**
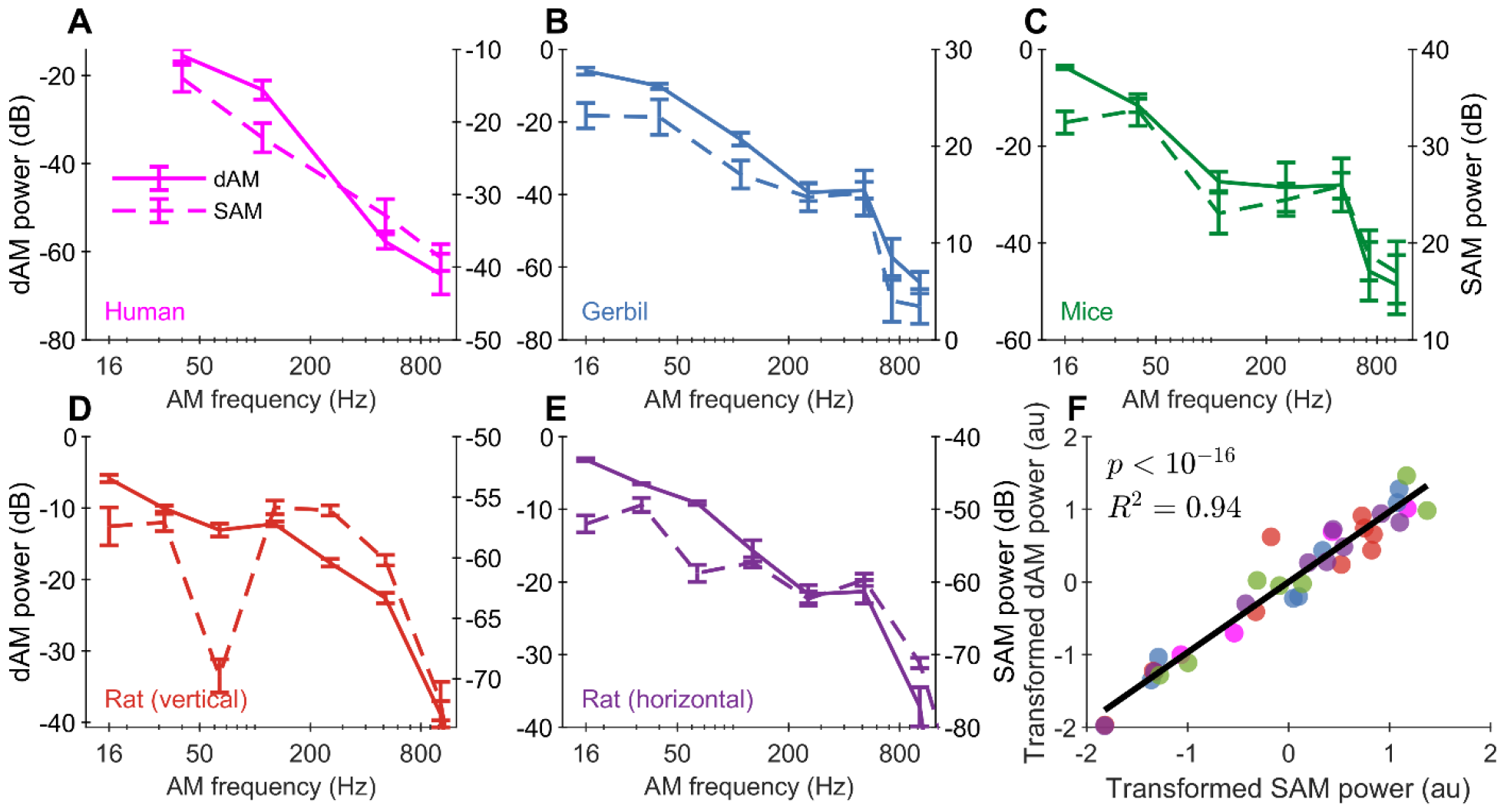
dAM-derived tMTFs are highly correlated with SAM-derived tMTFs. (A-E) Our dAM-derived tMTFs (left y-axis) and traditional SAM-derived tMTFs (right y-axis) for humans (A; *p* = .02, R^2^ = 0.94), gerbils (B; *p* = 1.5 × 10^−4^, R^2^ = 0.95), mice (C; *p* = 3.6 × 10^−4^, R^2^ = 0.92), and rats in vertical (D; *p* = 1.7 × 10^−4^, R^2^ = 0.87) and horizontal (E; *p* = 1.1 × 10^−6^, R^2^ = 0.97) recording configurations. (F) Scatter plot of transformed SAM tMTF power and transformed dAM tMTF power values for data in A-E. Data for each species and estimation method (i.e., SAM vs dAM) were transformed (normalized) to have zero mean and unity variance. Symbol colors correspond to species (same as in panels A-E).

## Discussion

Here, we present and validate a time-efficient method to estimate EFRs to AM frequencies, which has great clinical potential as an objective measure of the neural representation of speech-relevant temporal cues by the auditory system. Specifically, we introduce a dynamically varying AM tone or noise, which exponentially sweeps in AM frequency over a one-second duration (for e.g., from 16 Hz to 1500Hz, Fig. 1). Thus, the stimulus can capture AM coding by the entire neuraxis as different stations along the auditory pathway (e.g., frequencies below 80 Hz reflect cortical sources and frequencies above 300 Hz reflects subcortical sources). These responses can be analyzed using spectrally specific analyses, which offer spectrotemporally optimal resolution (Fig. 2). We show that high-SNR responses can be obtained up to 800 Hz in humans and 1.2 kHz in rodent species (Fig. 4). This approach can also be used to estimate the tMTF (Fig. 3), which are highly correlated with traditional tMTFs obtained using discrete and stationary AM tokens (Fig. 5).

Previous studies have focused on improving the efficiency of the auditory brainstem response, a tool primarily used to assess hearing thresholds in the clinic (Buran et al., 2020; Eysholdt & Schreiner, 1982; Polonenko & Maddox, 2019). However, to the best of our knowledge, a rapid frequency technique for EFRs to evaluate tracking temporal processing has not been published in the literature. While multiplexing-based approaches have also been employed to record envelope following responses (e.g., using multi-based harmonic complexes or multiple carrier-modulation combinations) (Holmes et al., 2018; John et al., 1998; Varghese et al., 2015; Wang et al., 2019), the stimulus we have employed is arguably more ecologically realistic than the typical stimuli (with static AM) employed in these studies, since naturally uttered speech, as well as animal vocalizations, consists of dynamic AM trajectories (Elliott & Theunissen, 2009; Singh & Theunissen, 2003). For example, the fundamental frequency contour can convey semantic (e.g., in Mandarin) as well as non-semantic (e.g., the emotional state of the speaker) information (Busso et al., 2009; Ververidis & Kotropoulos, 2006; Xu, 1997). Neural processing of these dynamic AM cues may be different depending on language/music experience (Chandrasekaran & Kraus, 2010; Krishnan et al., 2005; Wong et al., 2007), and could be more faithful in reflecting auditory processing of everyday sounds encountered by humans and animals. While here we show the proof of concept and validate the method for a single carrier (tone or noise), this method can be further optimized to parallelize the recording of EFRs for different frequency regions.

In addition to assessing dynamic neural tracking of arbitrary AM contours, the primary benefits of the proposed approach are its time efficiency and spectrotemporal resolution. For example, the traditional serial approach to estimate the tMTF (for five AM frequencies) requires ∼15 min of EFR data in our laboratories. In contrast, the EFR data to estimate the dAM-based tMTFs can be collected in ∼3 min, which offers a 5x time improvement. Furthermore, the recorded tMTFs are continuously sampled in time (and thus by design, in frequency because of the unique relationship between AM frequency and time). Therefore, as dAM trajectory sweeps across a wide range of frequencies, the tMTF can be estimated for any arbitrary AM frequency by choosing an appropriate window associated with that AM frequency. This is unlike the traditional SAM-based approach, where AM frequencies must be defined prior to data collection.

We have validated a rapid, cross-species method to evaluate temporal processing to use as a clinical and research tool for investigating suprathreshold deficits in the neural coding of speech-relevant temporal cues. This tool has the potential to be developed as an objective and rapid clinical measure of neural coding changes due to various hearing pathologies such as aging, noise exposure, dyslexia, autism spectrum disorders and auditory processing disorders (Bharadwaj et al., 2022; Maggu & Overath, 2021; McAnally & Stein, 1997; Parthasarathy & Kujawa, 2018; Shaheen et al., 2015), where the peripheral and central auditory pathways may be differentially affected.

## Methods

### Subjects

#### Humans

Young adult participants were recruited from the University of Pittsburgh Pitt + Me research participant registry, the University of Pittsburgh Department of Communication Science and Disorders research participant pool, and the community under a protocol approved by the University of Pittsburgh Institutional Review Board (IRB#21040125). Participants were compensated for their time and given an additional incentive to complete all study sessions. Participant eligibility was determined during the first session of the study. Eligible participants had normal cognition determined by the Montreal Cognitive Assessment (MoCA ≥ 25; Nasreddine et al., 2005), normal hearing thresholds (≤ 25 dB HL 250-8000 Hz), no severe tinnitus reported via the Tinnitus Handicap Inventory (THI; Newman et al., 1996), and Loudness Discomfort Levels (LDLs) > 80 dB HL at .5, 1, and 3kHz (Sherlock & Formby, 2005). Participants self-reported American English fluency. 13 participants (18-25 years old, 3 males) participants met these eligibility criteria and were tested further using the battery described below.

#### Gerbils and Mice

Fourteen Mongolian gerbils aged 18-27 weeks (male = 9) and six CBA/CaJ mice aged 52-57 weeks (male = 3) were used in this study. All animals are born and raised in our animal care facility from breeders obtained from Charles River (gerbils) or Jackson laboratories (mice). The acoustic environment within the holding facility has been characterized by noise-level data logging and is periodically monitored. Data logging revealed an average noise level of 56 dB, with transients not exceeding 74 dB during regular housing conditions and transients of 88dB once a week during cage changes. All animal procedures are approved by the Institutional Animal Care and Use Committee of the University of Pittsburgh (Protocol #21046600).

#### Rats

10 adult Fischer-344 rats aged 3-9 months obtained from Taconic were used in this study. The animals were housed in the animal care facility for the period of the study in relatively quiet, standard conditions. All protocols are approved by the Purdue animal care and use committee (PACUC 06-106).

### Experimental setup

#### Humans

Experiments were performed in a sound attenuating booth using a 64-channel EEG system (BioSemi ActiveTwo). Participants sat on a reclining chair and watched a silent, subtitled movie for the duration of the recording. Gold-foil tiptrodes were positioned in participants’ ear canals to deliver sound stimuli and record additional channels of evoked potentials from the ear canal. Stimuli were presented at 85 dB SPL with a carrier frequency of 3000 Hz. Averaged responses from the Fz to the ipsilateral tiptrode were further analyzed using custom-written scripts in MATLAB. Stimuli were generated using an Rx6 processor (TDT) and presented using ER3C insert earphones. Responses were recorded using the BioSemi software and additionally epoched and analyzed using external triggers that synchronized the stimulus and the response.

#### Animals

Experiments were performed in a double walled acoustic chamber. Animals were placed on a water circulated warming blanket set to 37 °C with the pump placed outside the recording chamber to eliminate audio and electrical interferences. Rats and gerbils were initially anesthetized with isoflurane gas anesthesia (4%) in an induction chamber. The animals were transferred post induction to a manifold and maintained at 1%–1.5% isoflurane. The electrodes were then positioned, and the animals were then injected with dexmedetomidine (Dexdomitor, 0.2-0.3 mg/kg intramuscularly for rats, 0.3 mg/kg subdermally for gerbils) and taken off the isoflurane. The usual duration of isoflurane anesthesia during this setup process was approximately 10 min. Recordings were commenced 15 min after cessation of isoflurane, with the time window for the effects of isoflurane to wear off determined empirically as 9 min, based on ABRs waveforms and latencies as well as the response to foot pinch stimuli. Dexmedetomidine is an alpha-adrenergic agonist which acts as a sedative and an analgesic and which is known to decrease motivation but preserve behavioral as well as neural responses in rodents (Ruotsalainen et al., 1997; Ter-Mikaelian et al., 2013). This helps to maintain animals in an un-anesthetized state, where they still respond to pain stimuli like a foot pinch but are otherwise compliant to recordings for a period of about 3 h. Mice were anesthetized with ketamine (100 mg/kg, i.p.) and xylazine (10 mg/kg, i.p.). Additional boosters at half the dose were given as necessary to maintain a stable place of anesthesia. The recordings were performed for mice under anesthetized conditions. The stimulus was presented to the right ear of the animal, freefield for rats (Bowers and Wilkins DM601 speaker) and mice (MF1 speakers, TDT), and using insert earphones (ER3C, Etymotic) for gerbils. Impedances from the electrodes were always less than 1 kHz as tested using the head-stage (RA4LI, Tucker Davis technologies, or TDT). Sounds were generated by SigGenRP (TDT). Signal presentation and acquisition was done by BioSig software (TDT) for rats, and a custom program for gerbils and mice (LabView). The output from the speakers and insert earphones was calibrated using a Bruel Kjaer microphone and SigCal (TDT) and was found to be within ±6 dB for the frequency range tested. Digitized waveforms were recorded with a multichannel recording and stimulation system (RZ-5, TDT) and analyzed with BioSig or custom written programs in MATLAB (Mathworks).

Subdermal (Ambu) were placed on the animals’ scalps for the recordings. In gerbils and mice, a positive electrode was placed along the vertex. For rats, a two-channel configuration was used similar to previous studies, with a positive electrode placed along the midline of the forehead for channel 1 (vertical configuration) and another positive electrode placed horizontally along the interaural line (horizontal configuration) for channel 2. The negative electrode was placed under the ipsilateral ear, along the mastoid, while the ground electrode was placed in the base of the tail.

### dAM Stimuli

FFRs were recorded in response to dynamically amplitude modulated stimuli with either a tone or noise carrier. For a desired AM frequency trajectory, *f*_*am*_(*t*), the phase trajectory [*φ*_*am*_(*t*)] can be estimated as its integration such that 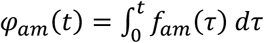 A dynamically modulated tone carrier (frequency *f* _*c*_, modulation depth *m*) can be obtained as 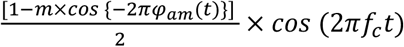. Similarly, a noise carrier, *η*(*t*), can be dynamically amplitude modulated as 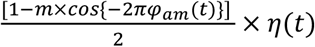

The tone carrier [*fc* = 3kHz (humans), 3 kHz (gerbils), and 12 kHz (mice)] consisted of two segments (each 500-ms long). In the first segment, *f*_*m*_ exponentially increased from 2.5 Hz to 40 Hz (frequency range assumed to receive substantial cortical contributions). In the second segment, *f*_*m*_ exponentially increased from 50 Hz to 1.2 kHz, which likely reflects subcortical contributions. An exponential increase was chosen to reflect the approximately logarithmic amplitude modulation processing by the auditory system (Ewert & Dau, 2000).

The noise carrier (used for rats) also consisted of two segments (each 500-ms long), where the first segment was modulated by AM exponentially increasing from 8 Hz to 120 Hz and the second segment was modulated by AM exponentially increasing from 45 Hz to 2.8 kHz. Both segments for the tone and noise carriers were played without any gap, which resulted in a slight click-like percept.

### Data analysis

EFRs were analyzed using traditional fast Fourier transforms (FFT) or recent spectrally specific analyses (Parida et al., 2021). For discrete EFRs, stimuli were presented in alternating polarity, with 500 repetitions in each polarity. Stimulus duration was 250 ms, and each AM token was presented at 3.1 repetitions/second, for a period of 322 ms. EFRs were sampled at a rate of 24414.1 Hz. EFRs were processed using a fourth-order Butterworth filter with a lowpass filter of 3000 Hz. The high-pass filter cutoffs used were 5 Hz, 80 Hz, 200 Hz, 300 Hz for 40 Hz, 110 Hz, 512 Hz, and 1024 Hz AM stimuli, respectively. Fast Fourier transforms (FFTs) were performed on the averaged time domain waveforms at each AM rate starting 10 ms after stimulus onset to exclude ABRs and ending 10ms after stimulus offset using MATLAB v. 2022a (MathWorks Inc., Natick, Massachusetts). The maximum amplitude of the FFT (Fast Fourier Transform) peak at one of three adjacent bins (∼3 Hz) around the modulation frequency of the AM rate is reported as the EFR amplitude.

More specific spectral metrics that capture the power along a known time-varying frequency trajectory (e.g., dynamic AM trajectory here) can be obtained using spectrally specific analyses comprising analytic signal extraction, frequency demodulation, and low-pass filtering. These analyses are particularly useful for neural data, like the FFR, which are stochastic and limited. Briefly, consider a signal, *x*(*t*), and a desired spectral trajectory, *f*_*am*_(*t*). First, the analytic signal can be estimated using the Hilbert transform. We define *a*(*t*) as 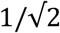 times the point-wise absolute magnitude of the analytic signal, where this normalization is needed to appropriately scale the power metrics defined next.

Next, the phase of the desired spectral trajectory can be obtained as 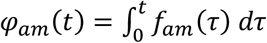.This phase signal is used to frequency demodulate *a*(*t*) as 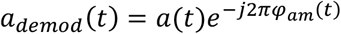 where power along *f*_*am*_(*t*) in *x*(*t*) [or *a*(*t*)] is mapped onto 0 Hz in *a*_*demod*_(*t*). Therefore, power along *fam*(*t*) in *x*(*t*) can be estimated as the power in *a*_*demod*_(*t*) in a low-frequency band centered around 0 Hz. To obtain this lowpass power metric, we computed a multitaper spectrum (using the Matlab pmtm function; time-bandwidth half product = 1.5) as it optimally minimizes bias and variance in the spectrum (Babadi & Brown, 2014; Thomson, 1982). Preliminary analysis showed that a 24 Hz (i.e., between -12 and 12 Hz) bandwidth for the low-pass filter offers sufficient resolution capturing most of the signal-driven EFR power with limited improvement with increasing bandwidth. We also estimated a noise floor using a parallel AM trajectory, which was shifted up by 36 Hz relative to the true dAM trajectory (to reduce overlap with it).

## Acknowledgements

This work was supported by the DOD HRRP RH200075 (ELB, AP), NIH-R21DC018882 (AP), NIDCD-R01DC013315 (trainee: SP), NIDCD T32DC011499 (Trainee: MEZ), and the PNC-Trees Charitable Trust (AP). We thank Megan Hallihan, Kathryn Bergstrom, Sarah Anthony, and Shaina Wasileski for their assistance with participant recruitment and data collection. The dynamic AM stimuli and spectrally specific analysis methods are currently patent pending (PCT International Application No. PCT/US2024/011312, claiming Priority from U.S. Provisional Patent Application No. 63/479,768).

